# Plasma proteomics reveals divergent sex-specific senescence and bone biology signatures across neurodegenerative diseases

**DOI:** 10.64898/2026.05.23.727339

**Authors:** Siddharth Kaipa, Ting Zhai, Sai Rama Krishna Meka, Patricia Moran-Losada, Zihuai He, Divya Channappa, Hamilton Se-Hwee Oh, Jarod Rutledge, Michael Zeineh, Anthony Agudelo, Ramesh Nair, Edward N. Wilson, Mounica Bandela, Keerthana Raghuraman, Amal Al-Rajhi, Anika Akkiraju, Janet Hwang, Frank M. Longo, Veronica Ramirez, Kathleen Poston, Nicola Neretti, Sundeep Khosla, MichaelP. Snyder, Tony Wyss-Coray, Victor W. Henderson, Vidyani Suryadevara

**Author notes:** Corresponding author: Vidyani Suryadevara, Department of Genetics, Stanford University, 240 Pasteur Dr Rm 4452, Palo Alto, CA 94304. equal contribution (Siddharth Kaipa, Ting Zhai).

## Abstract

Neurodegenerative diseases are often accompanied by systemic comorbidities, including changes in bone health, but the molecular relationship between neurodegeneration, skeletal decline, and cellular senescence remains poorly understood. In this study, we investigated sex-specific changes in circulating bone- and senescence-related proteins across the spectrum of Alzheimer’s disease(AD), Lewbody dementia(LB) and Parkinson’s disease(PD). Plasma proteomic profiling was performed on samples from 408 participants deeply phenotyped for neurodegenerative diseases, followed by differential protein and pathway analyses. This study reveals sex-dependent alterations in bone and senescence-related circulating proteins in AD-related, PD and LB-related neurodegenerative diseases, providing insights into the complex relationship between neurodegeneration and bone health. Several candidate proteins were also associated with established plasma neurodegeneration biomarkers, particularly pTau181. Pathway analyses revealed shared mitochondrial and metabolic dysfunction across neurodegenerative diseases, with disease-specific features including vesicle trafficking disruption in AD and inflammatory-senescence pathways in LB, plus sex-divergent patterns in inflammatory signaling and bone-related pathways.

## INTRODUCTION

Over 55 million people worldwide live with dementia, a number projected to rise to 139 million by 2050 ^1^, posing a significant socioeconomic burden; with women experiencing both higher prevalence and faster progressions^2^. One-third of adults globally have multiple chronic conditions, a risk that increases with age: two-thirds of those aged 65–85 and four-fifths of those over 85 have multiple conditions ^3^. Various multimorbidity patterns are linked to a higher dementia risk and contribute to mortality in affected patients ^4^. Studies indicate that sex affects both comorbidity profiles and the likelihood of developing Alzheimer’s disease (AD) and Alzheimer’s disease related dementias ^5^. The 2024 Lancet Commission estimated that at least 45% of dementia cases could be prevented, more than half of the attributable risk linked to hearing loss, elevated LDL cholesterol, hypertension, falls and fractures, diabetes, obesity, physical inactivity and smoking ^6^. These conditions impact multiple bodily systems, suggesting that various health issues may further heighten the risk of dementia. Nearly two-thirds of dementia cases are caused by AD ^7^, with other common causes being dementia with Lewy bodies (LB), vascular dementia, and frontotemporal dementia.

Emerging evidence shows sex-specific musculoskeletal decline in AD ^8,9^; consequently, falls and fractures are common, making bone-health management critical ^10^. Patients with early-stage AD show lower bone mineral density (BMD), which correlates with brain volume and cognitive decline ^11^. A year-long community-based prospective study from 1997-98 reported that individuals with mild cognitive impairment (Mini-Mental State Examination (MMSE) scores 18-23) had approximately twice the risk of developing hip fracture, and subjects with moderate-severe impairment (MMSE score < 18) had a similar relative risk ^12^. A prospective two-year cohort study from 2004 found that female AD participants with low bone mineral density (BMD) and low serum 25-hydroxyvitamin D concentrations with secondary hyperparathyroidism have an increased risk of hip fracture ^13^. A community-based prospective cohort study from 2005 showed that women, but not men, with low femoral neck BMD had approximately twice the incidence of AD and all-cause dementia ^14^. MMSE scores showed a significant association between cognitive impairment and low BMD in adults aged 50 years and above with no previous history of dementia or stroke, especially for women ^15^. A prospective five-year cohort study showed that low BMD and increased loss rate of BMD were associated with a higher risk of AD for both men and women ^14^. The most recent study shows a link between bone metabolic markers like serum osteocalcin and urine deoxypyridinoline/creatinine ratio and bone density in males with early-stage AD ^16^. Together, these findings highlight a connection between bone loss and cognitive decline, pointing to shared risk factors and aging mechanisms that affect both brain and bone health.

The common risk factors for dementia and impaired bone health are aging ^17^, *APOE* genotype ^18^, vitamin D deficiency, and lifestyle choices ^19^. Age is a primary risk factor for falls and fractures, as well as for AD, resulting in an increased accumulation of senescent cells with age, contributing to the pathogenesis of AD and many musculoskeletal conditions. In osteoarthritis (OA) and rheumatoid arthritis (RA), there are accumulations of senescent chondrocytes and senescent synoviocytes, causing senescence-associated secretory phenotype (SASP), disrupting joint homeostasis and resulting in cartilage damage ^20^. Senescence biomarkers, including SASP factors, have been reported in the neurons, microglia and astrocytes of AD brain tissues and cerebrospinal fluid (CSF) ^21^. Cellular mechanisms of AD involve the accumulation of amyloid-beta plaques and tau protein tangles, leading to synaptic dysfunction and neurodegeneration ^22^. Both aged wild-type mice and APP-PS1 mice (a preclinical model of AD) accumulate senescent bone marrow adipocytes and amyloid deposits, which can be cleared via senolytic treatment, preventing bone loss and clearing amyloidosis ^23^. Dasatinib and quercetin, navitoclax, and GW2580 among others have been promising senolytic treatments for the treatment of AD as well as musculoskeletal conditions ^24 25^. More broadly, senolytic therapies have been found to improve several diseases associated with aging including AD ^21^ and musculoskeletal diseases ^26^. This study may therefore help identify novel therapeutic strategies that simultaneously target bone and brain pathology.

Proteomics is a powerful tool for understanding the molecular complexity of the neurodegenerative diseases and cellular senescence, with strong potential for diagnostics, therapeutics, and personalized medicine. Unlike metabolomics and genomics, the proteomic approach offers a more accurate representation of a person’s current state, and the proteome tends to be more stable than the metabolome. This study is focused on identifying bone-associated changes in plasma of participants with different neurodegenerative disorders including AD spectrum (MCI, AD), and the LB spectrum (LB, MCI-PD) as well as Parkinson Disease (PD), as depicted in **Fig.1**. The objective of this study is to investigate sex-associated differences in bone and senescence markers of participants across the neurodegenerative spectrum and to provide mechanistic insights.

## METHODS

### Research participants

The Stanford Alzheimer’s Disease Research Center (SADRC) dataset contains 798 plasma samples from 408 individuals from a multi-ethnic volunteer cohort of older American adults (age 55-96, median 73) enrolled between 2015 and 2025 ^27^. This cohort is a longitudinal study of people with and without cognitive impairment, wherein some individuals have provided multiple plasma samples. Analyses are based on diagnostic groups: healthy controls (HC): without cognitive or motor impairment; AD spectrum: mild cognitive impairment (MCI) without motor impairment, and Alzheimer’s disease dementia; Lewy body (LB) spectrum: dementia with Lewy bodies and Parkinson’s disease dementia, mild cognitive impairment with Parkinson’s disease (MCI-PD) and Parkinson’s disease (PD) without cognitive impairment. All study protocols were approved by the Institutional Review Board at Stanford University. Participants or their legally authorized representatives provided written informed consent.

### Diagnosis of the AD spectrum

Clinical consensus diagnoses were based on published criteria for MCI ^28^, AD ^29^, dementia with Lewy bodies ^30^ and Parkinson’s disease dementia ^31^, MCI-PD ^32^, and Parkinson’s disease ^33^. Diagnoses were determined by a consensus panel that included neurologists, neuropsychologists, and research coordinators ^27^ based on neurological examination, neuropsychological testing, and functional assessment. Structural (MRI) and molecular (18F-florbetaben Aβ positron emission tomography [PET] and 18F-PI-2620 tau PET scan) neuroimaging was available for many participants. The Clinical Dementia Rating (CDR) obtained from a semi-structured interview with the participant and study partner served as a global indicator of cognition and function^34^. Participants provided blood specimens at each in-person visit. Participants diagnosed as HC (CDR = 0 or 0.5), MCI (CDR = 0, 0.5, or 1), or having AD dementia (CDR ≥ 0.5) were used in our analyses and treated as the discovery cohort. Cohort demographics and clinical diagnostic categories are summarized in **Suppl. Table 1**.

### Sample collection

To reduce variability due to blood draw and processing, a rigorous standardized protocol was followed in which around 10 cc of whole blood was collected in a vacutainer EDTA tube (Becton Dickinson) and centrifuged at 3,000 RPM for 10 minutes to separate the plasma. During plasma isolation, care was taken to leave 1 cm of plasma above the buffy coat while avoiding disturbance to the buffy coat to minimize cell contamination. The average plasma processing time, from blood draw to freezing and storage, was about 1 hour. All blood draws were performed in the morning after an overnight fast to reduce potential effects of diet and circadian variation on plasma protein levels.

### SOMAscan protein quantification and quality control

The SOMAscan assay platform (SomaLogic Inc.) was used to measure the relative levels of proteins in plasma samples. The plasma samples were analyzed using v.4.1 SomaScan assay ^35^. The platform is based on modified single-stranded DNA aptamers (SOMAmer) capable of binding to specific protein targets with high sensitivity and specificity. Plasma samples were analyzed in twelve different plates. To account for variation within and across batches, control, calibrator and buffer samples are added in each 96 well plate. The current analysis used 7,584 SOMAmer protein features after excluding 12 hybridization-control columns. Samples were restricted to SomaLogic RowCheck PASS sample rows; one QC-failed sample was excluded, and nine technical replicates were averaged on the log2 scale.

### Lumipulse plasma measurement

Lumipulse plasma measurements were conducted at the Stanford ADRC Biomarker Core and included pTau181, Aβ42 and Aβ40, and pTau217, GFAP, and NfL. Each sample was analyzed in singlicate per the automated platform’s high reproducibility as described previously ^36^. Briefly, plasma samples were thawed on wet ice, centrifuged for 5 min at 4°C at 1,000 × g, and loaded on a Lumipulse *G* 1200 instrument (Fujirebio US, Malvern, PA).

### Proteomics analysis

We analyzed the full proteome measured by SomaScan and two curated panels: 99 bone-related proteins represented by 133 SomaKeys, and 240 senescence-related proteins represented by 317 SomaKeys. We used the SenMayo ^37^ senescence marker panel to examine the changes in senescence-associated proteins in each of these neurodegenerative diseases. The list of bone and senescence markers used in this study are indicated in a table in Supplementary section (**Suppl. Table 2)**. Exact matches to the markers were identified in the data, and the somakeys were obtained.

We used linear mixed-effects models (LMMs) to evaluate differences in plasma protein abundance between each disease group and healthy controls. Protein abundances were modeled on the log2 RFU scale, with diagnostic group and age at visit as fixed effects and a participant-level random intercept to account for repeated samples from the same individual. In the combined-sex analysis, sex was included as an additional fixed effect: log2(Protein_ij) = β0 + γ0i + β1(Group_ij) + β2(Sex_ij) + β3(Age_ij) + ε_ij. Diagnostic groups were assigned at each visit, allowing participants whose diagnoses changed over time to contribute samples to the appropriate visit-level diagnostic group. Healthy controls were used as the reference group for all contrasts. Benjamini-Hochberg false discovery rate (FDR) correction was applied for multiple testing. In exploratory analyses, we used both nominal p<0.05 and within-panel FDR<0.2 for both relaxed significance cutoff and multiple-testing adjusted cutoff. Sex-stratified analyses were performed separately in males and females using the same pipeline, with sex omitted from the model: log2(Protein_ij) = β0 + γ0i + β1(Group_ij) + β2(Age_ij) + ε_ij.

### Pathway enrichment analysis

For each disease versus healthy-control comparison (combined and sex-stratified), proteins were ranked using a signed, scale-normalized statistic computed from the LMM fits, yielding one ranked list of approximately 6,329 ranked genes per comparison or stratum. SomaScan aptamers mapping to the same HGNC gene symbol were collapsed using the signed median z-statistic. Gene set enrichment analysis (GSEA) ^38^ was performed against the ranked lists using Reactome ^39^, MSigDB Hallmark ^40^, SenMayo ^37^, and our project-curated bone and senescence gene sets, with the Benjamini–Hochberg FDR computed within each stratum and pathway library. Senescence and bone biology were each evaluated against a small, hypothesis-aligned library of pathways grouped by mechanistic theme. The **senescence** library comprised: (i) the project-curated SenNet-derived senescence panel and the SenMayo signature as anchor references; (ii) five Reactome senescence sub-types;(iii) cell-cycle and DNA-damage-response pathways; (iv) anti-apoptotic / inflammatory SASP context; and (v) aging-context inflammation. The **bone** library comprised the project-curated bone panel together with Reactome bone-biology pathways and the Hallmark pathways included to capture sex-hormone–bone biology relevant to the sex-stratified design.

Aggregate enrichment scores can mask sub-components of a panel that move in opposite directions, so we performed a gene-level decomposition of each curated panel by hallmark sub-theme. Senescence genes were classified by the nine canonical SenNet hallmarks, with non-canonical members (keratin and cadherin tissue markers, skin-keratinization proteases, antiviral / interferon, vascular, WNT, MAPK, AD-specific) flagged separately so that tissue-of-origin or off-target contamination was not misread as senescence biology. Bone genes were similarly classified into bone-biology sub-themes spanning osteoblast / mineralization transcription, BMP, WNT, TGF-β and NOTCH signaling axes, bone resorption, calcium-phosphate endocrine regulation, collagens, MMP-mediated ECM remodeling, matricellular factors, IGFBPs / growth factors, and SASP-overlap inflammation. For each (sub-theme × disease × stratum) cell, we computed the arithmetic mean of z-statistics across the genes in that bin. This decomposition localizes the gene-level source of any panel-level signal, distinguishing, for example, signals driven by canonical senescence machinery (cell-cycle arrest, SASP secretome) from those driven by non-canonical tissue markers, and surfaces sex-divergent sub-theme patterns at finer resolution than the pathway-level statistics permit.

### Statistical analysis

To evaluate whether candidate plasma proteins identified in the disease-contrast analyses were associated with established markers of neurodegeneration, we tested their relationships with plasma pTau181, pTau217, GFAP, NfL, Aβ42, Aβ40, and Aβ42/40 ratio. For each biomarker-protein pair, we fit LMMs adjusted for baseline age, within-participant age change, sex, and individual diagnostic group, with a participant-level random intercept. Protein and biomarker values were modeled on the log2 scale; Aβ42/40 ratio was also evaluated on the raw ratio scale as a sensitivity analysis. Benjamini-Hochberg FDR correction was applied across all tested biomarker-protein associations within each candidate panel. Associations with FDR < 0.20 were considered statistically significant.

To characterize the proteomic landscape of neurodegeneration, differential protein expression was categorized by statistical stringency: nominally statistically significant proteins (p<0.05) are referred to as “nominal”, and FDR adjusted statistically significant proteins are referred to as “significant”. The labeled proteins highlight top 10 proteins with the smallest P values within each volcano plot. Some proteins might appear multiple times across these differential profiles because SomaScan platform measures some proteins utilizing multiple aptamers.

All statistical analyses were performed using R (version 4.4.1).

## RESULTS

### Alterations in bone-related proteins across neurodegeneration biomarkers

We assessed circulating bone-related proteins across the AD spectrum and subsequently performed sex-stratified analyses to evaluate molecular differences between females and males. The same analytical framework was applied to the LB spectrum and PD spectrum cohorts to assess whether bone and ECM-associated protein alterations differed across neurodegenerative diseases and by sex. Overall, our results indicate coordinated remodeling of extracellular matrix and bone-associated markers with sex-dependent differences in directionality and significance. In the unstratified AD spectrum cohort, IGFBP1 is nominally down regulated, and FN1 is significantly down regulated, while OLR1 is significantly up regulated **(Fig. 2A)**. Sex-stratfied analyses revealed distinct molecular dimorphisms. In females, the bone marker ANXA2 is nominally down regulated in the AD spectrum **(Supp.1A)**. In males, SERPINH1, COL2A1, FN1, and MMP14 are nominally down regulated, IGFBP1 is significantly down regulated, and OLR1 is significantly up regulated in the AD spectrum **(Supp.1D)**.

**Figure. 1:**
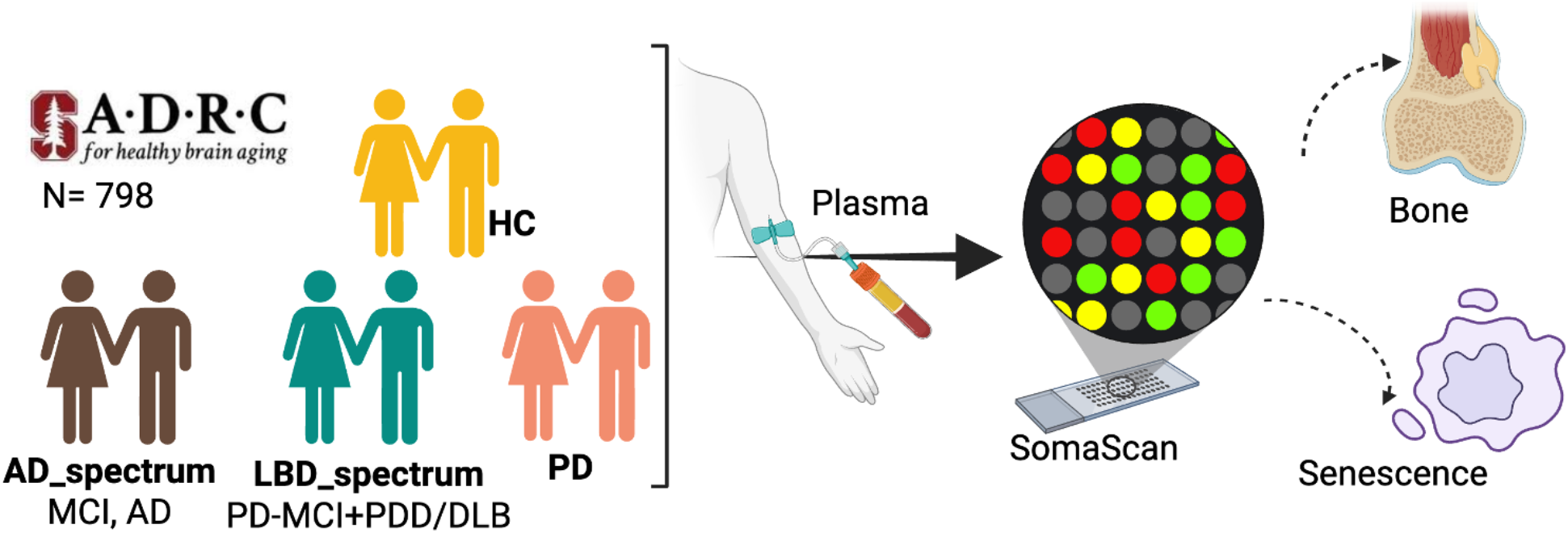
Experimental Design for Plasma Proteomic Profiling Across Neurodegenerative Disease Cohorts: Plasma samples were collected from healthy controls (HC) and participants with Alzheimer’s disease spectrum (AD_spectrum), Lewy body spectrum (LBD_spectrum), and Parkinson’s disease (PD) at Stanford Alzheimer’s Disease Center. Samples were analyzed using SomaScan aptamer-based proteomics platform, and proteomic signatures were correlated with bone metabolism and cellular senescence markers.

**Fig.2:**
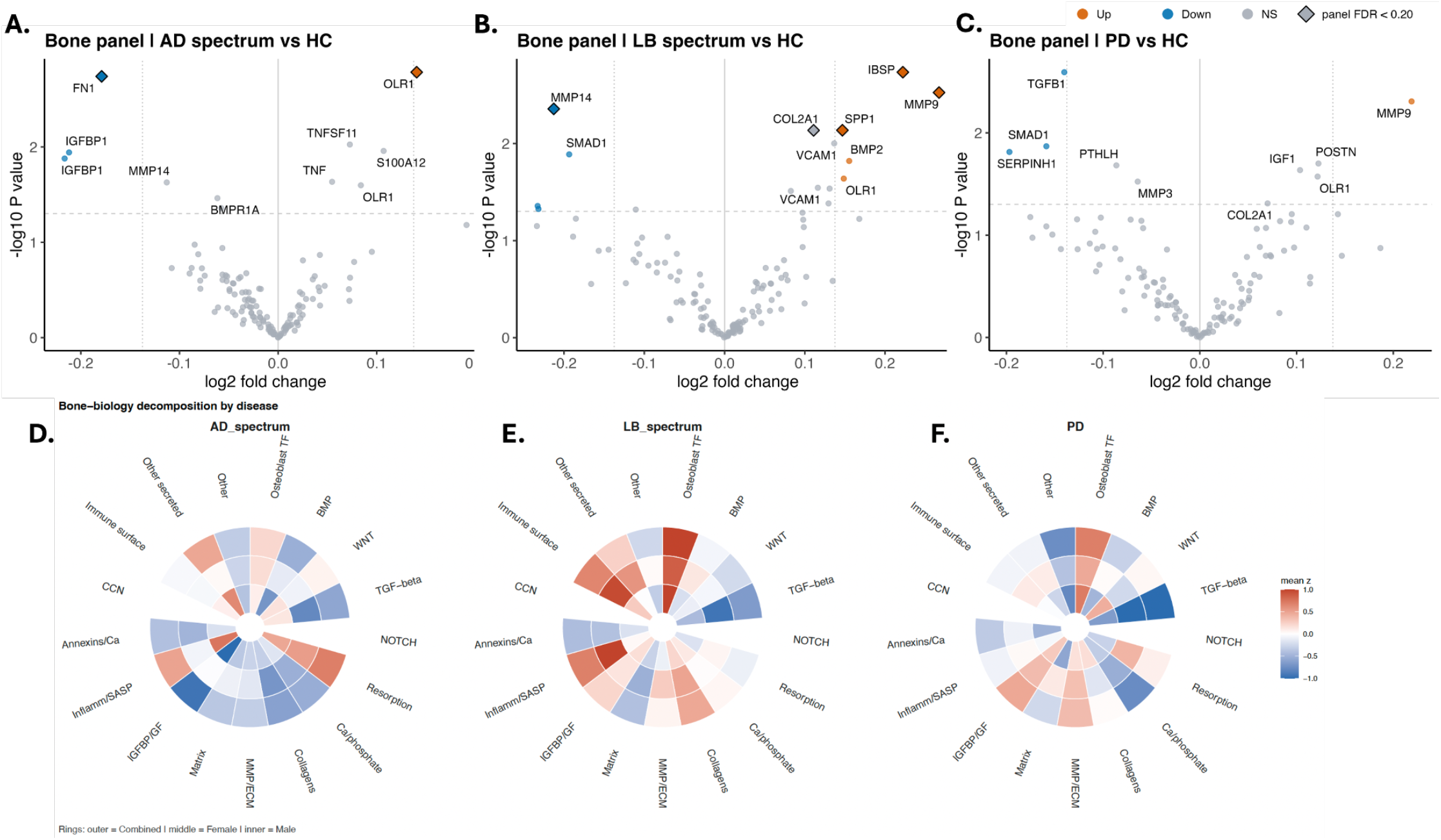
Bone-related proteins altered across cognitive impairment and neurodegenerative disease conditions: Volcano plots of differentially expressed bone-related proteins in plasma from patients with **A**. AD-spectrum, **B**. LB spectrum **C**. PD, compared with healthy controls (HC). Volcano plots illustrate the -log10 nominal p-value versus the log2 fold change in protein abundance relative to HC. A horizontal dashed line was drawn to indicate a nominal p-value of 0.05, and vertical dotted lines were drawn at ±0.138 log2 fold change, corresponding to 10% fold-change. Diamond markers indicate proteins passing panel-specific Benjamini-Hochberg FDR correction at FDR < 0.20. Top 10 markers are annotated. **Decomposition of the curated bone panel by bone-biology sub-theme:** Gene-level decomposition of the project-curated bone panel using the circular-rosette layout, with one rosette per disease **D**. AD-spectrum, **E**. LB-spectrum and **F**. PD and three concentric rings per panel (outermost = All, middle = Females, innermost = Males). Wedges correspond to bone-biology sub-themes defined by the analyst panel: osteoblast transcription factors and mineralisation; bone-morphogenetic-protein, WNT, TGF-β and NOTCH signalling axes; bone resorption and osteoclast biology; endocrine and calcium-phosphate regulation ; collagens and collagen processing; matrix-metalloprotease– mediated extracellular-matrix remodelling; matrix and matricellular factors; insulin-like growth-factor binding proteins and bone-relevant growth factors; SASP-overlap inflammation; annexin and Ca^2+^-dependent membrane proteins; the CCN matricellular family; bone-overlap immune-surface markers; and other bone-relevant secreted factors.

In the unstratified LB spectrum, SMAD1 is nominally down regulated, and MMP14 is significantly down regulated. Conversely, OLR1 and BMP2 are nominally up regulated, and SPP1, MMP9, IBSP, and COL2A1 are significantly up regulated **(Fig 2.B)**. When stratified by sex, profound differences emerged, in females, VCAM1, OLR1, and MEPE are nominally up regulated, and VCAM1, S100A12, MMP9, BMP2, IL1B, and COL10A1 are all significantly up regulated **(Supp.1B)**. In males, SMAD5, SMAD2, SERPINH1, and YWHAB are nominally down regulated, MMP14 and SMAD1 are significantly down regulated, and COL2A1, COL1A1, IBSP, and SPP1 are significantly up regulated **(Supp.1E)**.

Within the unstratified PD group, SERPINH1, SMAD1, and TGFB1 are nominally down regulated, whereas MMP9 is nominally upregulated **(Fig. 2C)**. Sex-specfic trends included: in females, TGFB1 is nominally down regulated, and OLR1 and MMP9 are nominally up regulated **(Supp.1C)**. Males demonstrated a broad downregulation, SERPINH1, SMAD5, YWHAB, and SMAD1 are significantly down regulated, CD69, SMAD2, PPP3CA, and PPP3R1 are nominally down regulated, while COL2A1 and COL1A1 are nominally up regulated (**Supp.1F**).

### Alterations in senescence-related proteins across neurodegeneration biomarkers

To further examine aging-related mechanisms underlying the bone–brain axis, we assessed circulating senescence-associated proteins across the neurodegenerative disease spectrum and evaluated sex-specific differences in senescence-related signaling. Given that cellular senescence is characterized by persistent cell-cycle arrest and the senescence-associated secretory phenotype (SASP), which promotes chronic inflammation, extracellular matrix remodeling, and altered tissue signaling, we focused on senescence linked cell-cycle regulatory networks and SASP-associated mediators. In the overall AD spectrum, the senescence-associated protein CDH12 is nominally down regulated **(Fig. 3A)**. VIM is nominally down regulated in females **(Supp.2A)**. In males, CDH12 and MMP14 are nominally down regulated, and IGFBP5 is significantly down regulated **(Supp.2D)**.

**Fig.3:**
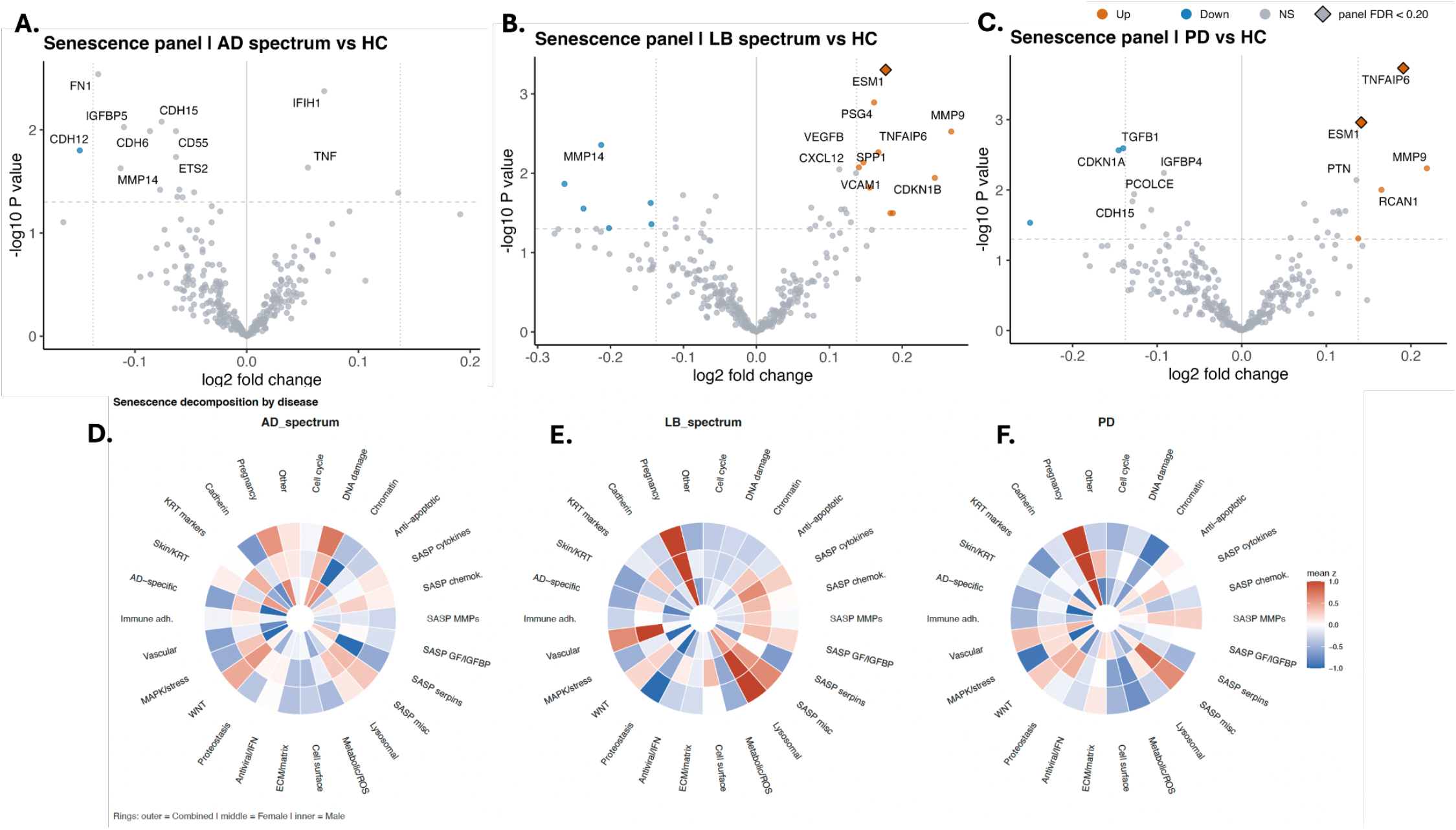
Senescence-related proteins altered across cognitive impairment and neurodegenerative disease cohorts: Volcano plots of differentially expressed bone-related proteins in plasma from patients with **A**. AD-spectrum, **B**. LB spectrum **C**. PD, compared with healthy controls (HC). Volcano plots illustrate the -log10 nominal p-value versus the log2 fold change in protein abundance relative to HC. A horizontal dashed line was drawn to indicate a nominal p-value of 0.05, and vertical dotted lines were drawn at ±0.138 log2 fold change, corresponding to 10% fold-change. Diamond markers indicate proteins passing panel-specific Benjamini-Hochberg FDR correction at FDR < 0.20. Top 10 markers are annotated. **Decomposition of the curated senescence panel by SenNet hallmark sub-theme:** Gene-level decomposition of the project-curated senescence panel, displayed as three circular rosettes **D**. AD-spectrum, **E**. LB-spectrum and **F**. PD. Within each panel, three concentric rings correspond to participant strata, with the outermost ring representing the combined cohort (All), the middle ring representing female participants (Females) and the innermost ring representing male participants (Males). Wedges around the perimeter correspond to sub-themes derived from the nine canonical hallmarks of the SenNet framework together with non-canonical components flagged separately to prevent tissue-of-origin or off-target signals from being misread as senescence. Wedge fill encodes the arithmetic mean of the LMM-derived z-statistic across the genes assigned to each sub-theme on a divergent red–blue scale (red, up-regulated in disease; blue, down-regulated; midpoint at zero).

Across the unstratified LB spectrum, ESM1 is significantly up-regulated, MMP14 is nominally down regulated in the LB spectrum, while CDKN1B, VCAM1, CXCL12, SPP1, TNFAIP6, MMP9, VEGFB, and PSG4 are nominally up regulated **(Fig. 3B)**. Sex-stratified analyses revealed pronounced differences. Female participants showed a coordinated up-regulation signature, with significant elevation of ESM1, HGF, IL1B, TNFAIP6, BMP2, PTGS2, TEK, MMP9, VCAM1, PAPPA, and RTP4 (**Supp.2B**). In contrast, males displayed a divergent pattern characterized by nominal down-regulation of VSIR, APOBEC3G, FGF7, and IGFBP4, together with significant down-regulation of MMP14. These decreases occurred alongside nominal up-regulation of CCL3, VEGFB, and CDKN1B, and significant up-regulation of SPP1 (**Supp.2E**).

Within the PD cohort, CDKN1A and TGFB1 are nominally down regulated, whereas RCAN1 and MMP9 are nominally up regulated. Concurrently, ESM1 and TNFAIP6 are significantly up regulated **(Fig 3.C)**. In females, TGFB1 is nominally down regulated, and PCOLCE is significantly down regulated. IL15 and PAPPA are nominally up regulated, and TNFAIP6, MMP9, RCAN1, and ESM1 are all significantly up regulated (**Supp.2C)**. Males demonstrated a predominantly downregulatory trend including CXCL3, MAPK3, CDKN1A, KRT1, CDKN2D, MAPRE1, and CLIC4, In contrast, CRIM1 was significantly upregulated, while a few targets including PSG5 were nominally upregulated within this group **(Supp.2F)**.

### Validation of bone and senescence markers with conventional neurodegeneration biomarkers

To evaluate whether the candidate plasma proteins identified through bone and senescence in the above analyses relate to established markers of neurodegeneration, we examined/tested their associations with conventional plasma biomarkers (pTau181, pTau217, GFAP, NfL, AB142, AB140, and AB ratio), using 18 bone- and 17 senescence-panel candidate proteins discovered at panel FDR<0.20 in the three disease-vs-control contrasts (AD spectrum, LB spectrum and PD). 131 bone–biomarker associations across all 18 candidates and 114 senescence–biomarker associations across all 17 candidates reached FDR<0.20 (**Fig 4**), 94 and 79 survived FDR<0.10. pTau181 contributed the most hits in both panels. Among bone candidates, MMP-14, IGFBP-1, IBSP and COL10A1 were inversely associated with pTau181 (β = −0.09 to −0.22), while YWHAB (14-3-3 β/α), SMAD5, SERPINH1 and SMAD1 were positively associated (β up to +0.53). Among senescence candidates, MMP-14, TSG-6 (TNFAIP6), Endocan (ESM1), sTie-2 (TEK) and CRIM1 were inversely related to pTau181, while RTP4 and PTGS2 were positively related. One of the largest inverse biomarker associations was between MMP-9 and the Aβ42/40 ratio in females. Significant associations were more numerous in females than males in the bone panel, while senescence-panel associations were broadly similar between sexes.

**Fig.4.**
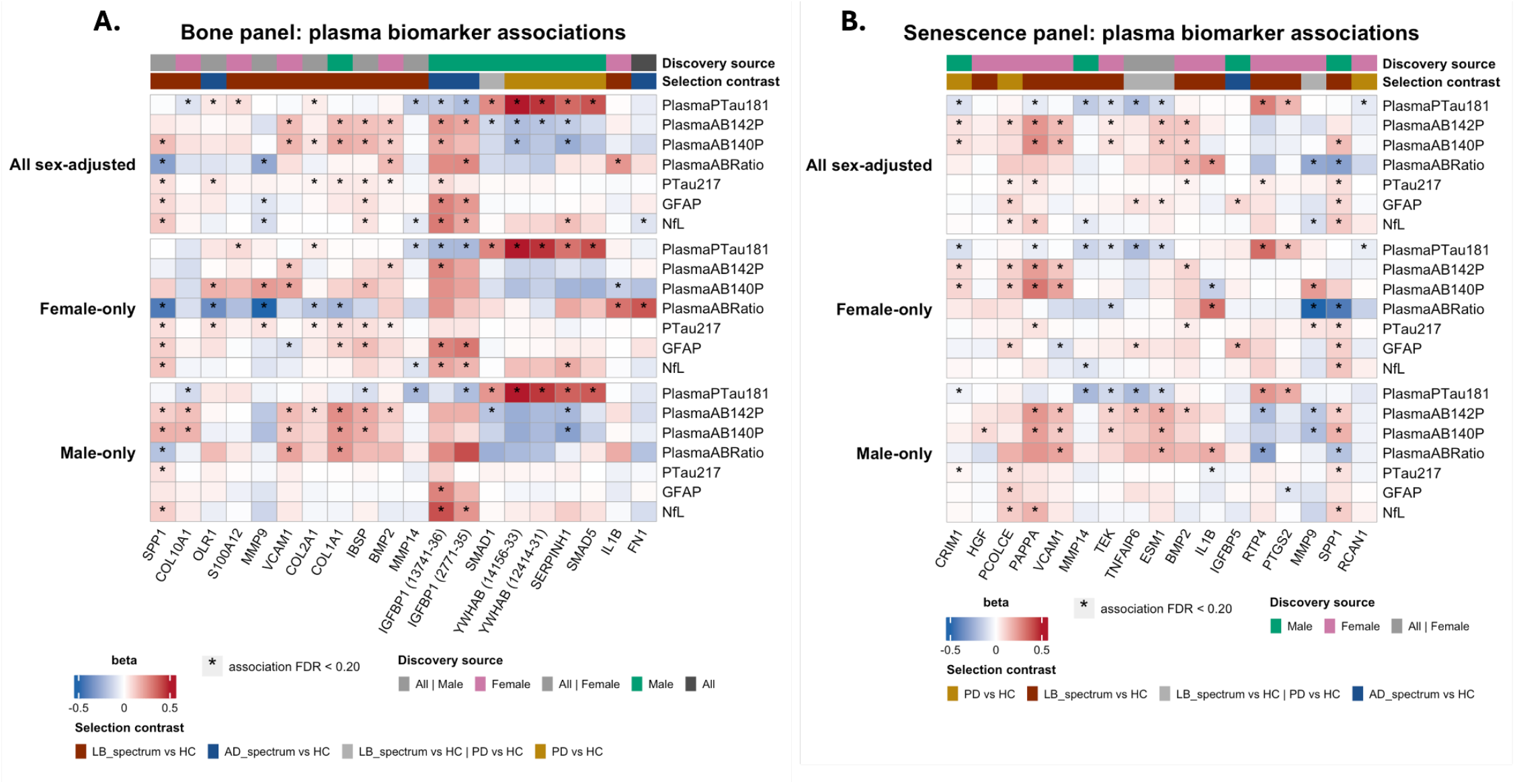
Validation of candidate plasma protein signals and conventional neurodegeneration biomarkers: **A**. Bone-related candidate proteins. **B**. Senescence-related candidate proteins. Heatmaps display regression coefficients from biomarker–protein association models for the same selected protein set across three analysis strata: all participants with sex adjustment, female-only, and male-only. Models included a random intercept for participants to account for repeated visits and adjusted for age and diagnosis group; the all-participant model additionally adjusted for sex. Protein values were analyzed on the log2 RFU scale, and biomarker values were log2-transformed before modeling. Candidate proteins were selected from individual diagnostic discovery contrasts using discovery FDR < 0.20. The top annotation bars indicate the discovery source and selection contrast for each protein. Asterisks denote biomarker–protein associations passing Benjamini–Hochberg FDR correction within each analysis stratum (FDR<0.20). Color scales represent regression coefficients (red = positive association; blue = negative association).

**Fig.5.**
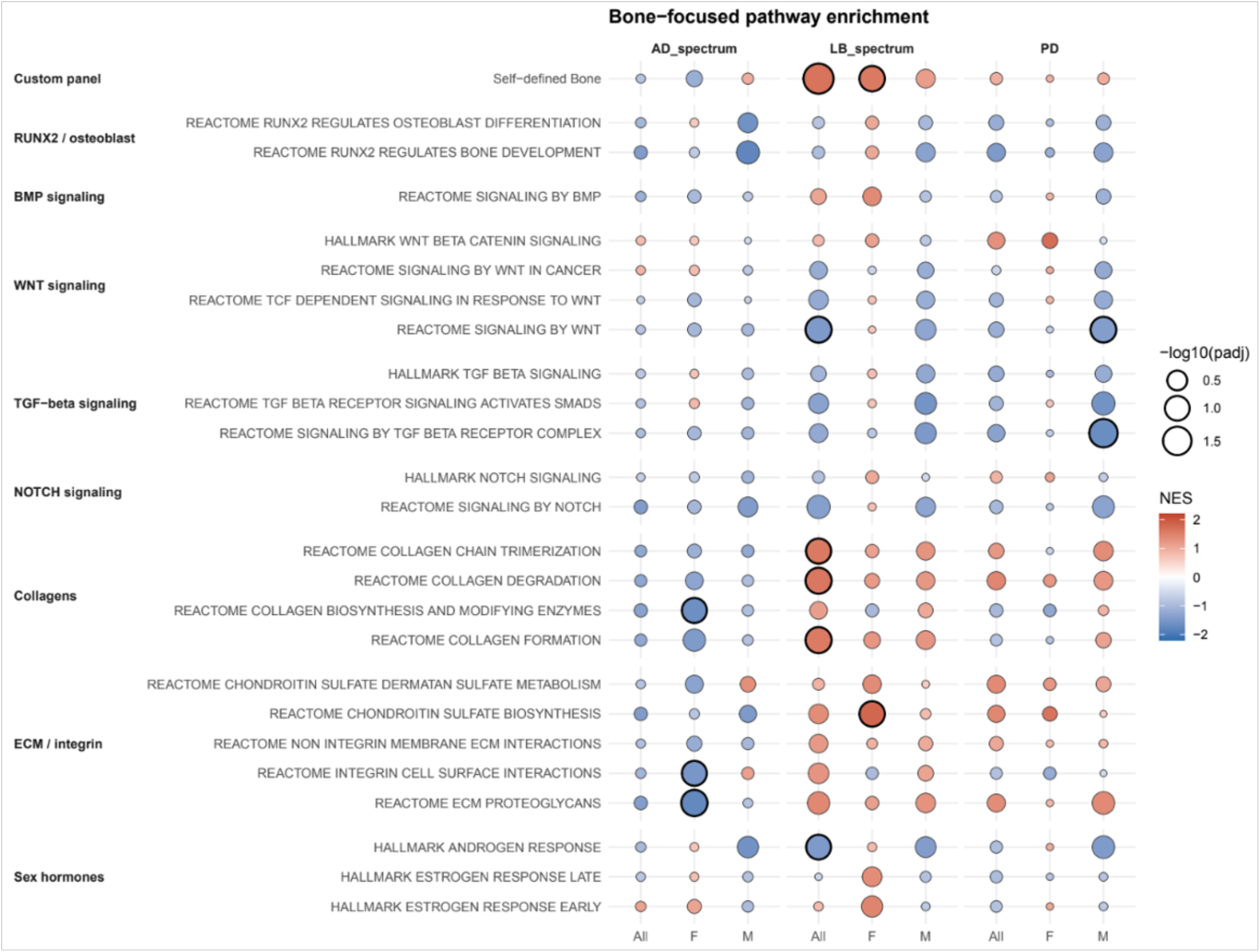
Bone-focused pathway enrichment across neurodegenerative diseases: Preranked gene set enrichment analysis of plasma proteins against a targeted bone-biology pathway library. Rows show the project-curated bone panel together with Reactome and Hallmark pathways grouped by bone-biology theme: RUNX2-regulated bone development and osteoblast differentiation (Reactome); BMP signalling (Reactome); WNT signalling (Reactome and Hallmark); TGF-β signalling (Reactome and Hallmark); NOTCH signalling (Reactome and Hallmark); collagen biosynthesis, chain trimerisation, and degradation (Reactome); extracellular matrix, chondroitin-sulfate metabolism, and integrin interactions (Reactome); and sex-hormone responses (Hallmark estrogen-response early/late and androgen response), included to capture the sex-hormone–bone axis relevant to the sex-stratified design. Columns show disease contrasts grouped by AD-spectrum, LB-spectrum, and PD, with All, female-only, and male-only strata within each contrast. Dot fill encodes the normalised enrichment score (NES) on a divergent red–blue scale (red, up-regulated in disease; blue, down-regulated). Dot size encodes−log_10_(adjusted p-value), with Benjamini–Hochberg false-discovery-rate correction applied within each stratum across the targeted library.

**Fig.6.**
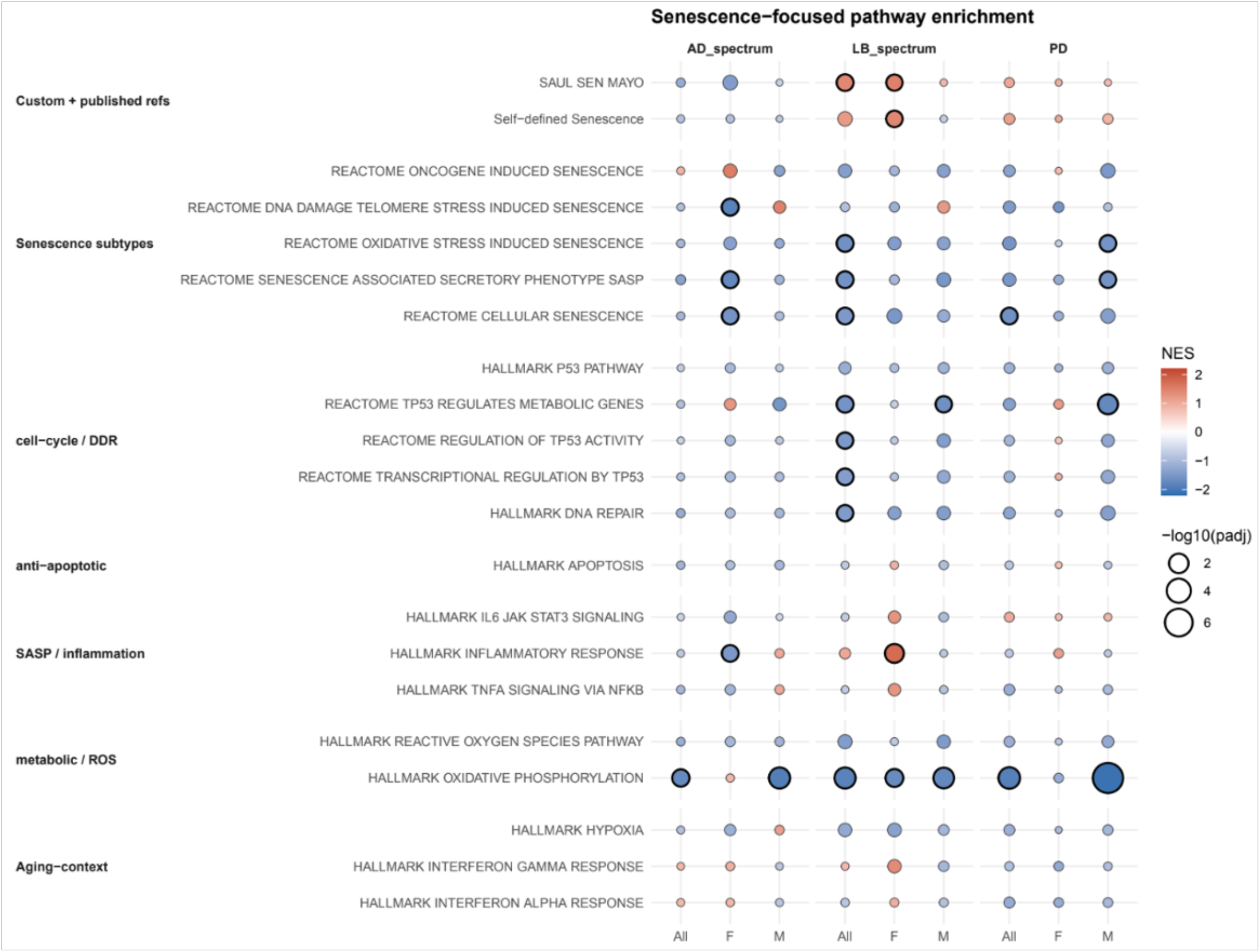
Senescence-focused pathway enrichment across neurodegenerative diseases: Preranked gene set enrichment analysis of plasma proteins ranked by their linear mixed-effects model z-statistic for each disease-versus-healthy-control comparison. Rows show pathways from a targeted senescence library, grouped by mechanistic theme: the project-curated SenNet-derived senescence panel and the published SenMayo signature as anchor references; five Reactome senescence subtypes; cell-cycle and DNA-damage-response programs; the Hallmark apoptosis set; SASP-aligned inflammatory pathways; metabolic and reactive-oxygen-species pathways; and aging-context pathways. Columns show disease contrasts grouped by AD-spectrum, LB-spectrum, and PD, with All, female-only, and male-only strata within each contrast. Dot fill encodes the normalised enrichment score (NES) on a divergent red–blue scale (red, up-regulated in disease; blue, down-regulated). Dot size encodes −log_10_(adjusted p-value), with Benjamini–Hochberg false-discovery-rate correction applied within each stratum across the targeted library. SASP, senescence-associated secretory phenotype.

### Enrichment analysis reveals shared and disease-specific molecular pathways across neurodegenerative diseases

The bone panel reached significance in the LB spectrum, including the whole-cohort contrast and female participants. In the whole LB spectrum, this was accompanied by up-regulation of extracellular-matrix organisation and collagen formation/degradation pathways, whereas in LB-female participants the strongest bone-axis signals included the curated bone panel and chondroitin-sulfate biosynthesis (**Fig.5**). A reciprocal, female-specific pattern appeared in the AD spectrum, where extracellular-matrix organisation, collagen biosynthesis, ECM proteoglycans and integrin–cell-surface interactions were significantly down-regulated in females. Other bone-axis pathways were male-restricted: TGF-β and WNT signalling were down-regulated in PD-male participants, and Hallmark androgen response was down-regulated in the whole LB comparison.

Mitochondrial oxidative phosphorylation was significantly down-regulated in every disease comparison: AD spectrum, LB spectrum and PD vs healthy controls, and in male participants of each, identifying a shared peripheral metabolic-senescence signal across the neurodegenerative spectrum (**Fig.6**). The LB spectrum carried the broadest senescence-pathway disruption. When the entire LB group was compared with healthy controls, cellular senescence, oxidative-stress-induced senescence, the SASP gene set, DNA repair and TP53 transcriptional/metabolic programmes were all significantly down-regulated, while the published SenMayo senescence signature was significantly up-regulated, a Lewy-body–specific elevation absent in AD and PD. In female participants with LB spectrum, Hallmark inflammatory response, the project-curated SenNet senescence panel and SenMayo were coherently up-regulated. Male LB participants showed disruption of the oxidative-phosphorylation / MYC-targets / TP53-metabolic axis.

Two sex-restricted senescence signals stood out beyond LB. Female participants with AD spectrum showed significant down-regulation of telomere maintenance, the DNA-damage / telomere-stress response, the SASP gene set, cellular senescence and Hallmark inflammatory response, whereas AD-male participants were limited to the oxidative-phosphorylation signal. Male PD participants additionally down-regulated the SASP gene set and oxidative-stress-induced senescence, showing that a SASP-suppressed pattern is not unique to AD-female.

Gene-level decomposition clarified these aggregate findings. The bone rosettes (**Fig. 2D–F**) showed heterogeneous bone-axis remodeling across disease groups and strata. Bone-resorption / osteoclast markers were most consistently positive in the AD spectrum, but showed mixed patterns in the LB spectrum and PD group. IGFBP / growth-factor, TGF-β, and WNT-related sub-themes showed sex- and disease-dependent directionality rather than uniform up- or down-regulation. Osteoblast-transcription / mineralisation markers showed a modest, mixed pattern and did not indicate a clear AD-male-specific deficit. The senescence rosettes (**Fig.3 D–F**) showed coherent SASP-secretome up-regulation in LB participants (MMPs/TIMPs, growth factors, IGFBPs, chemokines); concurrent down-regulation of cell-cycle and SASP machinery against up-regulation of keratin, WNT and AD-associated genes (APP, APOE) in AD-female participants, explaining the modest aggregate NES; and a PD pattern dominated by down-regulated cell-cycle and DNA-damage-response sub-themes.

## DISCUSSION

Proteomics is essential for studying disease pathophysiology because proteins drive almost all biological processes ^41^. The field is particularly crucial for mapping the mechanisms behind complex disorders like neurodegenerative diseases, their comorbidities and aging-related processes such as cellular senescence ^42^. Plasma proteomics, in particular, offers powerful insights into disease mechanisms, biomarker discovery, and therapeutic development. In this study, we phenotyped participants across neurodegenerative diseases against healthy controls to investigate the relationship between bone-brain health, with the goal of identifying proteomic markers for bone and senescence proteins, which link brain-bone health deterioration and to determine whether these markers have sex-dependent variation.

Neurodegenerative disease, such as Alzheimer’s Disease, is associated with comorbidities such as periodontitis ^43^, depression ^44^, diabetes ^45^, chronic kidney disease ^46^, inflammatory bowel disease ^47^, and cerebrovascular disease ^48^. The intersection of brain and bone pathology is highly recognized, with preclinical models offering mechanistic insights and therapeutic strategies into this relationship. In 3-month-old 3xTg-AD mice, diminished bone quality and less fracture resistance were observed, which was found to be reversed by treatment with resveratrol and exercise ^49^. Similarly, when mice with markedly reduced bone mass were given lithium, their bone metabolism and bone mass were restored to near wild-type levels ^50^, while chronic lithium treatment was found to reduce amyloid-β production and senile plaque formation, leading to an improvement in spatial learning and memory abilities ^51^. Another novel therapy that involves inhibition of FSH using an anti-FSHβ antibody inhibits plaque and neurofibrillary tangle formation and reverses cognitive decline in ovariectomized 3xTg-AD mice ^52^, while a humanized version of the anti-FSH antibody called MS-Hu6 showed an increase of bone formation in the femur and spine of humanized Tg32 mice ^53^. Collectively, these observations aim at identifying therapeutic targets that can simultaneously target brain and bone pathologies.

Our proteomic analyses revealed significant alterations in bone-related proteins across the neurodegenerative spectrum, with several notable sex-dependent differences. Particularly, the elevation of matrix remodeling enzymes such as MMP-9 aligns with the effects of a pro-inflammatory skeletal profile. In female participants specifically, bone-related proteins involved in extracellular matrix remodeling (MMP-9), osteoclastic / mineralisation biology (BMP-2, COL10A1, S100A12) and adhesion (VCAM-1) were significantly elevated, with additional nominal trends in WNT signalling (WNT16) and BMP antagonism (GREM1); these female-specific signals were concentrated in the LB spectrum. The detailed functions of all the significant proteins in bone and brain are provided in **Table 2**. The bone related proteins that are altered in the plasma of patients with neurodegenerative disease included in this study have been implicated in various bone diseases including osteoarthritis and osteoporosis. Among proteins altered in our cohort that have been previously implicated in osteoporosis are IGFBP-1, FGF-19, BMP-2, SMAD1/5, MEPE, collagen-binding protein (SERPINH1), WNT16, GREM1, BSP (IBSP), osteopontin (SPP1), COL10A1, S100A12 and IL-1β. 10-year long epidemiological evidence from Korea indicated that osteoporosis contributed to an increased risk for dementia ^54^. Further, patients with osteoporosis or fragility fractures treated with nitrogen-containing bisphosphonates had a lower risk for dementia ^55^ and studies have shown that AD patients have increased risk for fractures than those without AD ^56^, indicating a bi-directional implication of AD and osteoporosis. The connection between neurodegeneration and bone health extends beyond osteoporosis to osteoarthritis. Meta-analysis and data from UK biobank have shown that osteoarthritis is linked to increased risk for dementia ^57^ and mouse model studies of AD have found that knee osteoarthritis induced by destabilizing the medial meniscus in 5xFAD mice results in accelerated amyloid plaque deposition and neurodegeneration ^58^. Several proteins attributed to osteoarthritis were also altered, including MMP-9, MMP-14, chondrocalcin (COL2A1 C-propeptide), BMP-2, S100A12, IL-1β, WNT16, GREM1, VCAM-1, osteopontin (SPP1) and collagen-binding protein (SERPINH1), further emphasizing the molecular crosstalk between skeletal and neurological disease processes.

At the pathway level, our analysis revealed a peripheral metabolic-senescence signature shared across all three disease spectra (down-regulated mitochondrial oxidative phosphorylation), against which two disease- and sex-specific patterns were superimposed. The Lewy-body spectrum showed the broadest senescence-pathway disruption, coordinate down-regulation of cellular senescence, SASP and DNA-damage / TP53 programmes, accompanied by up-regulation of the published SenMayo signature and, in female participants, Hallmark inflammatory response. This female-skewed inflammatory–senescence axis in the Lewy-body spectrum is consistent with post-mortem evidence of neuroinflammation and innate immune activation in dementia with Lewy bodies ^59^, and with a possible peripheral interferon-response component captured by the RTP4 signal in our candidate set ^60^. The down-regulation of p53-regulated metabolic programmes that we observed in male PD and male Lewy-body participants is in line with the previously reported role of p53 in dopaminergic neuron vulnerability ^61^. In contrast, female AD participants displayed a coherent suppression of SASP, telomere-stress and ECM-organisation programmes, while the project-curated bone panel reached significance only in the Lewy-body spectrum, with up-regulated collagen and extracellular-matrix remodelling, a directionally opposite signal to the ECM/collagen down-regulation seen in female AD. Together, these findings suggest that the bone–brain axis reflects both shared and disease-specific biology, with conserved peripheral metabolic and bone-resorption signatures across neurodegenerative diseases superimposed on sex-dependent inflammatory and extracellular matrix remodeling programs, particularly the contrasting ECM/SASP elevation in female Lewy-body disease and suppression in female AD.

Given that neurodegeneration and bone deterioration are both fundamentally age-associated pathophysiologies, we further investigated senescence-related proteins, and identified significant alterations in all participants, with notable sex-specific differences. Several senescence-associated secretory phenotype (SASP) factors, TSG-6, Endocan (ESM1), HGF, IL-1β, BMP-2, MMP-9, MCP-3 (CCL7), RTP4, PAPP-A, VCAM-1, sTie-2 (TEK) and PTGS2 (COX-2), were significantly up-regulated in the Lewy-body spectrum, with the strongest signal in female participants, while the AD spectrum showed predominantly down-regulated IGFBP-family senescence markers (IGFBP-5, IGFBP-6) in male participants. These findings suggest that senescence-associated pathways play a crucial role in the pathogenesis of neurodegenerative diseases as previously reported ^62^, and that sex-specific mechanisms may influence the accumulation of senescent cells and their impact on disease progression. The circulating biomarkers detected in this study are distinct from each other, thus revealing various patterns of manifestation of the circulating markers and identifying bone- and senescent-proteins that are more closely related to the changes in brain health. The proteins detected in the plasma indicate high abundance proteins produced throughout the body including those involved in inflammation, coagulation and metabolism, which collectively underscore the systemic nature of neurodegenerative disease.

### Limitations of the study

A key limitation of this study is the moderate cohort size (n = 408); while this provides a solid foundation, future research should aim to include a larger and more demographically diverse sample size to validate our findings and enhance the robustness of our identifications. In the future, we plan to perform similar proteomic analysis from additional large cohorts, and also analyze approximately 40,000 samples harmonized across 22 different centers within the Global Proteomics consortium. A notable strength of this ADRC cohort is the well-characterized neurodegenerative pathologies encompassing imaging, biomarker and cognitive assessments. These findings will be a foundation for future mechanistic studies focusing on elucidating the molecular mechanisms underlying these sex-specific alterations and their impact on disease progression. Additionally, longitudinal studies are required to determine the temporal dynamics of these biomarkers and their potential as predictive indicators of disease severity and patient outcomes.

In conclusion, this study reveals sex-dependent alterations in bone and senescence-related circulating proteins in AD-related and in LB-related neurodegenerative diseases, providing insights into the complex relationship between neurodegeneration and bone health. These findings suggest that sex-specific therapeutic strategies may be necessary to effectively manage bone-related comorbidities in people with dementia and underscore the importance of adopting a holistic approach to neurodegenerative disease management, considering bone health, senescence, and biological sex-specific factors during aging.

## ACKNOWLEDGEMENTS

This work was supported by the Stanford Alzheimer’s Disease Research Center (National Institute on Aging grants P50AG047366 and P30AG066515), the National Institute on Aging (AG072255,T.W.-C; AG057909, AG061155 and AG044829, S.M. and N.B; AG066206, Z.H.), the National Institutes of Health (R01AG044546, RF1AG053303, RF1AG058501 and U01AG058922, C.C.; P01AG003991, C.C. and J.C.M.; RF1AG074007,Y.J.S.), the Michael J. Fox Foundation (L.I. and C.C.), the Alzheimer’s Association Zenith Fellows Award (ZEN-22-848604, C.C.), the Milky Way Research Foundation, Nan Fung Life Sciences (T.W.-C.), the Stanford Graduate Fellowship (H.O. and J.R.), the Stanford Translational Program in Aging Research (T32AG0471). We would like to thank Phil and Penny Knight Initiative for Brain Resilience for funding the measurement of AD fluid biomarkers. We are grateful to the Jackson Lab tissue mapping center at the NIH SenNet consortium for allowing us to use their group’s senescence gene list. We thank the participants whose generosity enabled this study.

## AUTHOR CONTRIBUTIONS

Conceptualization, V.S., V.H.; methodology, V.S., T.Z., V.H., S.K., T.W.C; Somascan platform proteomics: J.R., H.O., V.R. and P.M.-L; Statistical methods: N.N., Z.H. and T.Z.; Plasma collection led by D.C with K.R., A.AR.; AD biomarkers: E.W.; Senescence expertise: N.N.; Bone expertise: Su.K.; Overall insights: M.Z., M.S.; V.W.H. assisted in Stanford-ADRC data acquisition. Stanford ADRC led by V.W.H., F.M.L. and T.W.-C.; Figures: S.K., V.S., T.Z, formal analysis, S.K., T.Z.; writing: S.K., V.S., T.Z, M.B. supervision, V.S., V.H.;

## DATA AND CODE AVAILABILITY

Stanford-ADRC data are available upon reasonable request to the Stanford-ADRC data release committee, https://web.stanford.edu/group/adrc/cgi-bin/web-proj/datareq.php. All analysis scripts are publicly available through github repository: https://github.com/zhaiting/ADRC-proteomics.

