## Supplementary material for "Plasma proteomics reveals divergent sex-specific senescence and bone biology signatures across neurodegenerative diseases": NA

**Suppl. Table 1:** Demographics and disease state of Stanford ADRC participants from whom plasma were collected.

| **Characteristic** | **HC** | **AD spectrum** | **PD** | **LB spectrum** |
| --- | --- | --- | --- | --- |
| Model sample rows, n | 380 | 217 | 124 | 77 |
| Unique participants, n | 197 | 120 | 73 | 46 |
| Age, years at first included visit in group, median [IQR] | 73 [68, 77] | 76 [68, 80] | 70 [65, 74] | 72 [64, 78] |
| Female participants, n (%) | 122 (61.9%) | 56 (46.7%) | 34 (46.6%) | 14 (30.4%) |
| Included visits per participant, median [range] | 2 [1-4] | 1 [1-5] | 2 [1-3] | 1 [1-4] |
| Participants with one included visit, n (%) | 65 (33.0%) | 61 (50.8%) | 30 (41.1%) | 24 (52.2%) |

Cohort construction: From the 830-sample QC-passed SomaScan set, 9 technical replicates were averaged on log2 scale; samples with baseline age <55 years were excluded; 3 participants flagged in clinical notes as unconfirmed MCI were dropped (4 sample-visits across 3 subjects). The final analytical set consisted of 798 sample-visits and 408 participants. Diagnostic group columns reflect each participant's visits while in that group; they are not mutually exclusive because 34 participants contribute visits to more than one diagnostic group during follow-up. Participant-level age and sex summaries use each participant's first included visit within the displayed group; the Overall column uses each participant's first included visit overall. 3 participants have sex labels that vary across visits; the first-included-visit rule resolves them for participant-level summaries.

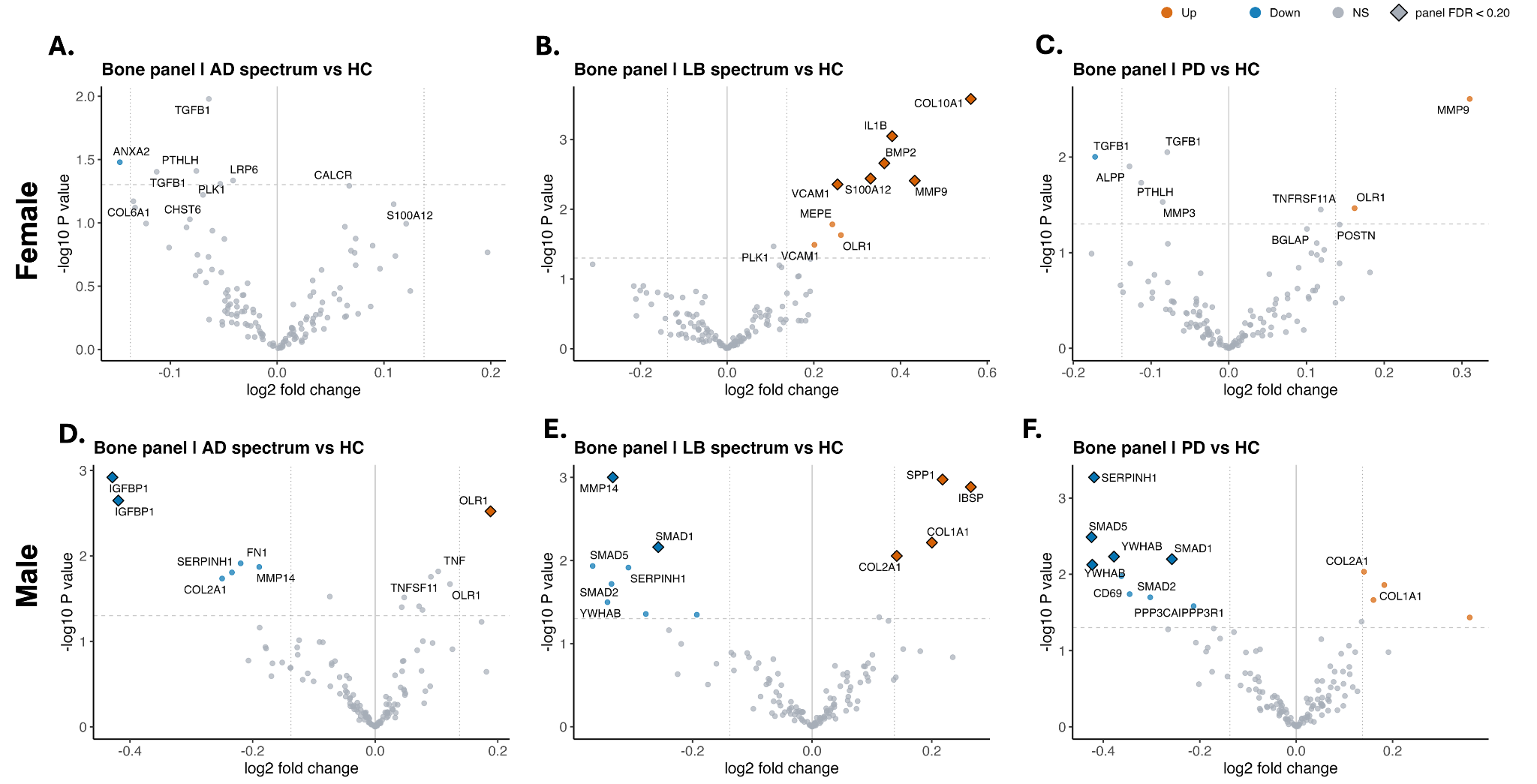
**Suppl. Fig.1: Sex-specific alterations in bone-related proteins across neurodegenerative disease cohorts:** Sex-stratified volcano plots of differentially expressed bone-related proteins versus HC. Females **(A–C)** and males **(D–F)** are shown across AD **(A,D)**, LB **(B,E)**, and PD **(C,F)** spectra. Colored dots represent proteins that are suggestive discovery (nominal p value < 0.05), while proteins not meeting this threshold are in gray. Volcano plots illustrate the -log10 nominal p-value versus the log2 fold change in protein abundance relative to HC. A horizontal dashed line was drawn to indicate a nominal p-value of 0.05, and vertical dotted lines were drawn at ±0.138 log2 fold change, corresponding to 10% fold-change.

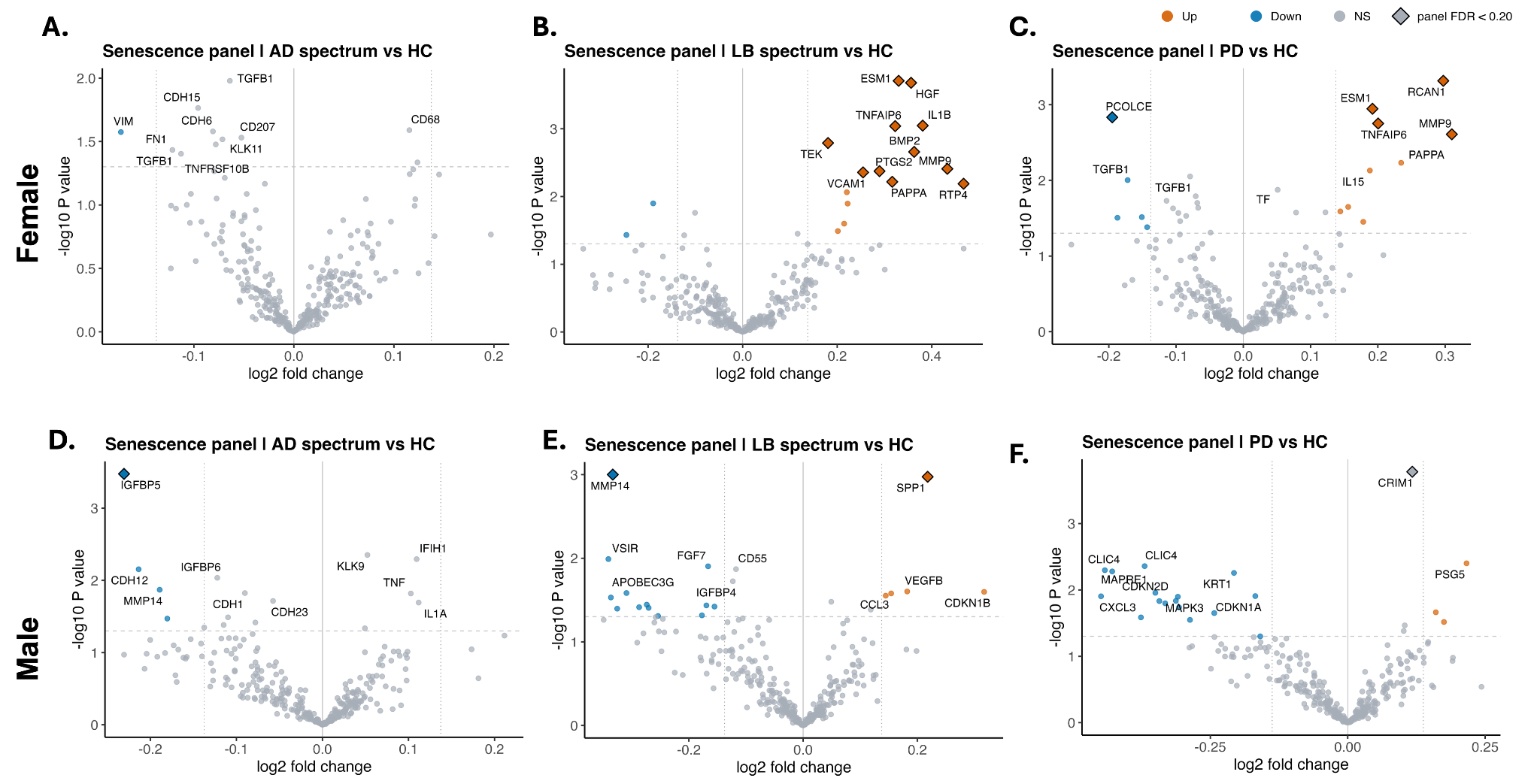

**Suppl. Fig.2. Senescence-related proteins altered in the plasma of neurodegenerative disease cohorts:** Sex-stratified volcano plots of differentially expressed senescence-related proteins versus HC. Females **(A–C)** and males **(D–F)** are shown across AD **(A,D)**, LB **(B,E)**, and PD **(C,F)** spectra. Colored dots represent proteins that are suggestive discovery (nominal p value < 0.05), while proteins not meeting this threshold are shown in gray. Volcano plots illustrate the -log10 nominal p-value versus the log2 fold change in protein abundance relative to HC. A horizontal dashed line was drawn to indicate a nominal p-value of 0.05, and vertical dotted lines were drawn at ±0.138 log2 fold change, corresponding to 10% fold-change.

| **Bone** | ALP; BGLAP; COL1A1; COL1A2; RUNX2; SPP1; BMP2; BMP7; DMP1; MEPE; PHEX; CTSK; MMP9; TRAP; TNFSF11; TNFRSF11B; MMP13; CALCR; PTH; FGF23; TGFB1; LRP5; LRP6; SOST; VCAM1; CSF1; ANXA2; ANXA5; TNF; IL6; COL10A1; COL3A1; VEGFA; TGFBR1; OPG; IGF1; PDGF; GREM1; NOG; ACVR1; FBN1; SOX9; SPARC; ZFYVE21; VAMP4; TNFSF8; TMEM100; TNS3; TNN; CCT8; PLK1; CD163; OLAH; RGN; S100A12; PTHD3; PSCA; PROM1; MKI67; POSTN; SLC26A4; OLR1; NBR1; ND4; ND2; ND5; ND3; SIRT7; STEAP3; MMP9; KDM5D; LGR6; ITGB3; ITGAV; IRAG2; IGSF2; GPC3; LGALS10; ENPP1; CD69; UHRF1; ADAM19; COX3; TNF-a; CO1; CYTB; CDA; CCNDBP1; GCM2; CEP170; CEACAM8; CD48; CLEC12A; JMJD6; BSG; AHSP; ATP6; ARID5B; YWHAB; COL2A1; COL10A1; COL3A1; IBSP; SPARC; MGP; MMP2; RANKL; Osteoprotegerin; RUNX2; Sclerostin; Cathepsin K; Calcitonin; Integrin Beta-3; RANK; Vascular Cell Adhesion Molecule-1; Insulin-Like Growth Factor 1; Platelet-Derived Growth Factor; Fibronectin; Vitronectin; Transforming Growth Factor Beta; Bone Alkaline Phosphatase; Vitamin D; Estrogen; Testosterone; Glucocorticoids; Tartrate-Resistant Acid Phosphatase; Sirtuins; SIRT7; Dickkopf-1; ANXA2; ANXA5; Matrix Metalloproteinase 3; MMP3; Alkaline Phosphatase; Placental Type; ALPP; Calcitonin Gene-Related Peptide; CGRP; Parathyroid Hormone-Related Protein; PTHrP; Fibroblast Growth Factor 2; FGF2; Interleukin 1 Beta; IL-1β; IL-1B; Interleukin 17; IL-17; C-terminal Telopeptide of Type I Collagen; CTX-I; N-terminal Propeptide of Type I Collagen; P1NP; TRANCE; RANKL; TRAP; Bone Morphogenetic Protein 3; BMP3; Bone Morphogenetic Protein 4 ; BMP4; Calcium-sensing Receptor; CaSR; Calcineurin; SUMF1; Collagen a1(VI); CHST6; IGFBP-1; Chondrocalcin; FGF-19; WNT16; MMP-1; Collagen-binding protein; YKL-40; MMP20; MMP14; PLOD2; Thrombospondin-1; Ephrin-A2; FZD9; BMP-1; BMP-4; BMPR1A; BSP; SUMF1; MMP-1; SMAD1; SMAD2; SMAD5; WISP-1; WNT5A; WNT5B |
| --- | --- |
| **Senescence** | CDKN2A; p16INK4A; p16; CDKN1A; p21CIP1; p21; IL-6; TNF; H2AX; IL-1α; IL-1a; IL-1β; IL-1b; SA-β-gal; SA-B-gal; SERPINE; PAI-1; TGFβ1; TGFB1; p53; CCL2; CCL5; CXCL1; CXCL8; MMP12; CCL3; HMGB1; MMP2; CCL8; CXCL10; CXCL2; GDF15; IGFBP3; IGFBP-3; MMP3; MMP9; CSF2; IGFBP7; IGFBP-7; LMNB1; MMP1; BCL2; CCL4; CCL7; CDKN2B; p15INK4B; p15; CSF1; CXCL12; FAS; ICAM1; IFNγ; IFNy; IGF1; IGFBP4; IGFBP-4; IL-17A; MAPK14; MMP13; NFKB1; Telomere length; TIMP2; TNFRSF1B; VEGFA; CSF3; CXCL16; HGF; ICAM3; IGFBP2; IGFBP-2; IGFBP5; IGFBP-5; IL-7; INHBA; Lipofuscin; MKI67; MMP10; PLAUR; SPP1; TAF; TNFRSF1A; TP53BP1; ACVR1B; ANG; ANGPT1; ANGPTL4; AREG; AXL; BEX3; BMP2; BMP6; C3; CCL1; CCL13; CCL16; CCL2; CCL20; CCL24; CCL26; CCL3; CCL3L1; CCL4; CCL5; CCL7; CCL8; CD55; CD9; CSF1; CSF2; CSF2RB; CST4; CTNNB1; CTSB; CXCL1; CXCL10; CXCL12; CXCL16; CXCL2; CXCL3; CXCL8; CXCR2; DKK1; EDN1; EGF; EGFR; EREG; ESM1; ETS2; FAS; FGF1; FGF2; FGF7; GDF15; GEM; GMFG; HGF; HMGB1; ICAM1; ICAM3; IGF1; IGFBP1; IGFBP2; IGFBP3; IGFBP4; IGFBP5; IGFBP6; IGFBP7; IL10; IL13; IL15; IL18; IL1A; IL1B; IL2; IL32; IL6; IL6ST; IL7; INHA; IQGAP2; ITGA2; ITPKA; JUN; KITLG; LCP1; MIF; MMP1; MMP10; MMP12; MMP13; MMP14; MMP2; MMP3; MMP9; NAP1L4; NRG1; PAPPA; PECAM1; PGF; PIGF; PLAT; PLAU; PLAUR; PTBP1; PTGER2; PTGES; RPS6KA5; SCAMP4; SELPLG; SEMA3F; SERPINB4; SERPINE1; SERPINE2; SPP1; SPX; TIMP2; TNF; TNFRSF10C; TNFRSF11B; TNFRSF1A; TNFRSF1B; TUBGCP2; VEGFA; VEGFC; VGF; WNT16; WNT2 |

**Suppl. Table 2:** List of bone and senescent related markers
